# RoDiCE: Robust differential protein co-expression analysis for cancer complexome

**DOI:** 10.1101/2020.12.22.423973

**Authors:** Yusuke Matsui, Yuichi Abe, Kohei Uno, Satoru Miyano

## Abstract

**Motivation:** The full spectrum of abnormalities in cancer-associated protein complexes remains largely unknown. Comparing the co-expression structure of each protein complex between tumor and healthy cells may provide insights regarding cancer-specific protein dysfunction. However, the technical limitations of mass spectrometry-based proteomics, including contamination with biological protein variants, causes noise that leads to non-negligible over- (or under-) estimating co-expression.

**Results:** We propose a robust algorithm for identifying protein complex aberrations in cancer based on differential protein co-expression testing. Our method based on a copula is sufficient for improving identification accuracy with noisy data compared to conventional linear correlation-based approaches. As an application, we use large-scale proteomic data from renal cancer to show that important protein complexes, regulatory signaling pathways, and drug targets can be identified. The proposed approach surpasses traditional linear correlations to provide insights into higher order differential co-expression structures.

**Availability and Implementation:** https://github.com/ymatts/RoDiCE.

## 1 Introduction

Cancer is a complex system driven by many molecular events, including genomic mutations, as well as epigenetic and transcriptomic dysregulation (Hoadley et al., 2018). However, our knowledge regarding how their upstream events characterize downstream mechanisms with proteomic phenotypes remains scarce (Clark et al., 2019; Liu et al., 2016; Mertins et al., 2016; Zhang et al., 2016). Protein complexes are responsible for most cellular activities. In fact, recent studies (Ori et al., 2016; Romanov et al., 2019; Ryan et al., 2017) have demonstrated that protein subunits tend to exhibit co-expression patterns in proteome profiles. Furthermore, protein complex subunits are simultaneously down- or up-regulated via genomic mutations (Ryan et al., 2017). However, little is known regarding the changes that occur in the co-regulation of protein complexes between tumor and normal healthy tissues.

Accordingly, in the current study, we propose a novel algorithm for differential co-expression of protein abundance to identify tumor-specific abnormalities in protein complexes. Differential co-expression (DC) analysis is a standard technique for gene expression analysis to identify differential modes of co-regulation between conditions. As such, numerous methods already exist (Bhuva et al., 2019), including correlation analysis, which is one of the most common measures of co-expression. For example, differential correlation analysis (DiffCorr) (Fukushima, 2013) and gene set co-expression analysis (GSCA) (Choi and Kendziorski, 2009) are two-sample Pearson’s correlation coefficient tests. However, studies report that protein expression levels have greater variability than gene expression levels due to post-translational modification regulatory mechanisms (Gunawardana et al., 2015; Liu et al., 2016). This variability can affect the estimation of co-expression as an outlier and can significantly impact DC results.

We, therefore, developed a robust DC framework, designated robust differential co-expression analysis (RoDiCE), via two-sample randomization tests with empirical copula. The notable advantage of RoDiCE is noise robustness. Our main contributions are as follows: 1) development of an efficient algorithm for robust copula-based statistical DC testing; 2) overcome computational hurdles associated with the copula-based permutation test by incorporating extreme value theory; 3) demonstrate the effective application of copula to cancer complexome analysis; and 4) develop a computationally efficient multi-thread implemented as an R package.

### 1.1 Motivational example from the CPTAC/TCGA dataset

First, to demonstrate the need for robustness in protein co-expression analysis, we analyzed a cancer proteome dataset of clear renal cell carcinoma from CPTAC/TCGA with 110 tumor tissue samples. We measured co-expression using Pearson’s correlation coefficient and compared the correlation coefficients before and after removing outliers. To identify outlier samples, we applied robust principal component analysis using the R package ROBPCA (Hubert et al., 2005) with default parameters. Among 49,635,666 pairs of 9,964 proteins, the correlation coefficients of 7,541,853 (15.2%) pairs were deviated by more than 0.2 after removing outlier samples (**Figure 1**). Note, the value “0.2” is provided as an example of the number of pairs with a change in correlation coefficient, and does not, therefore, have statistical or biological meaning. This result implied that a non-negligible proportion of protein co-expression would be overestimated or underestimated. To accurately compare the structures of co-expression, it is necessary to compare them while minimizing co-expression over-/under-estimation.

**Fig 1.**
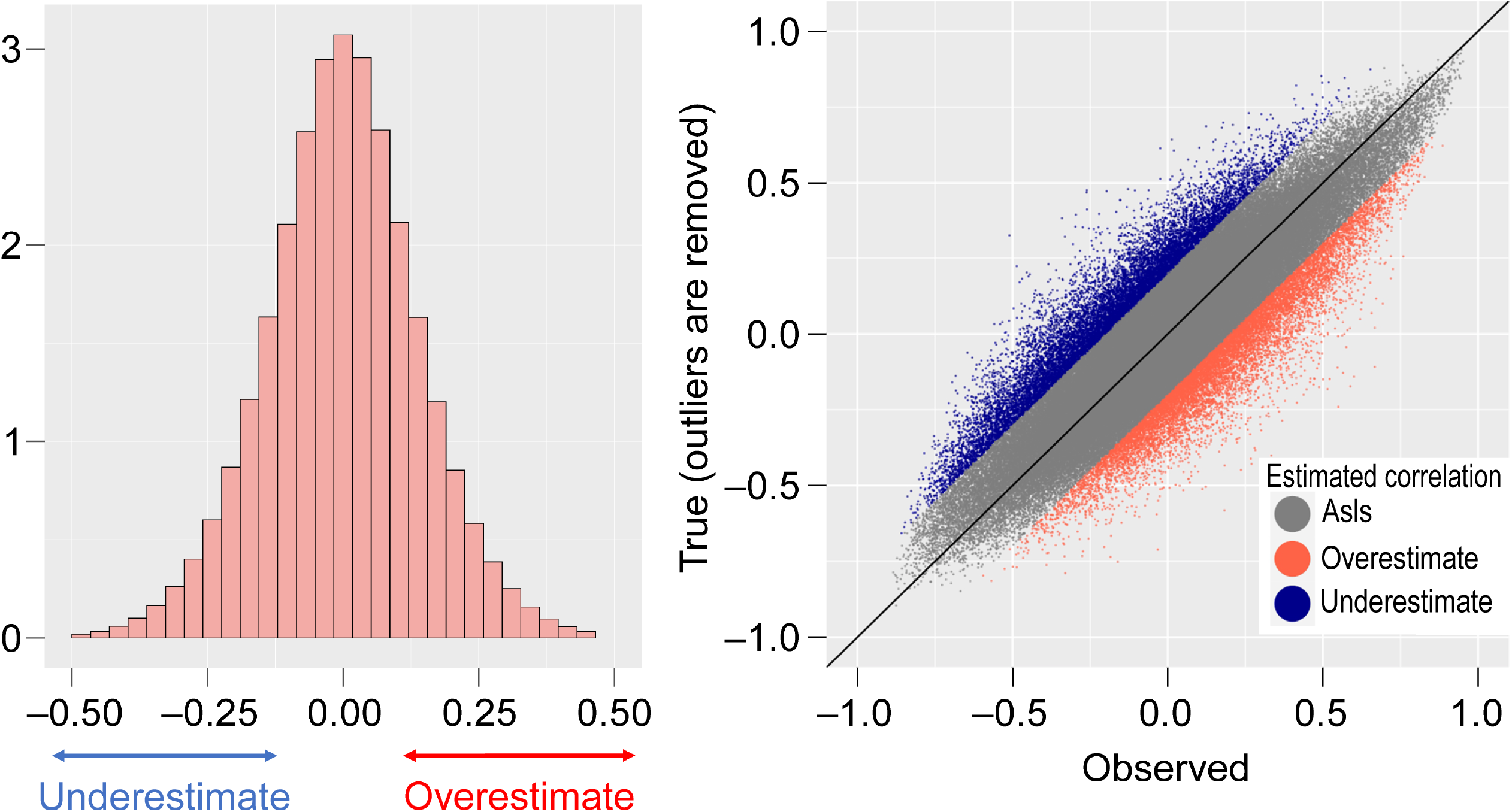
Example of outlier effects on co-expression. Difference in Pearson’s correlation before and after removing outlier samples; the left panel shows a histogram of the difference in correlation differences. The right panel shows a scatter plot of the original correlation against one without outlier samples.

## 2 Methods

Figure 2 provides an overview of RoDiCE. We decomposed the expression level of subunits in the protein complex into a structure representing co-expression and one representing the expression level of each subunit, using the empirical copula function (Nelsen, 2010). The empirical copula rank converts the scale of the original data. By comparing the empirical copula functions with the conditions for statistical hypothesis testing, we derived the *p*-value as the differences in co-expression structures. Our method is described in detail in the following sections.

**Fig 2.**
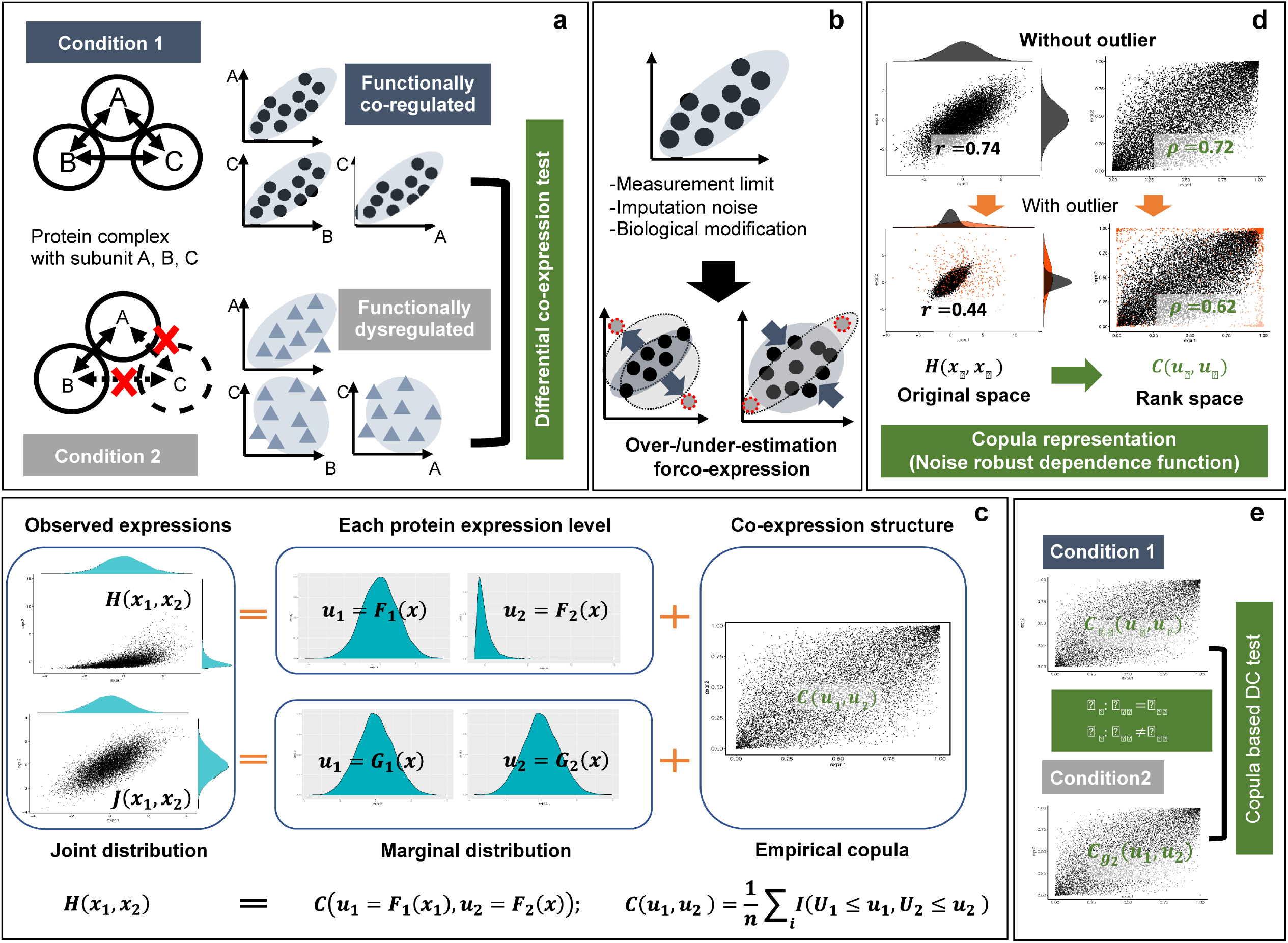
Overview of RoDiCE. **a) Objective of analysis via RoDiCE.** The proposed method aims to identify abnormal protein complexes by comparing two abnormal groups. An abnormal complex is one where the co-expressed structure is different in at least two subunits. **b) Protein co-expression and outliers.** The protein expression levels measured through LC/MS/MS contain some outliers due to the addition of noise from several sources. These can cause over- (or under-) estimation in the co-expression structure. **c) Copula decomposition.** The RoDiCE model decomposes the observed joint distribution of protein expression into a marginal distribution representing the behavior of each protein and an empirical copula function representing the latent co-expression structures between proteins. This allows for the extraction of potential co-expressed structures and for their robust comparison against outliers. The figure shows an example where the co-expressed structure estimated by the copula is the same for two apparently different joint distributions of protein expression. **d) Copula robustness**. A copula is a function that expresses a dependency on a rank-transformed space of data scales. One advantage of transforming the original scale into a space of rank scale is that it is robust to outliers. The example in the figure compares Pearson’s linear correlations with Pearson’s linear correlations in the space converted to a rank scale by a copula function (Spearman’s linear correlations). Pearson’s linear correlation underestimates from 0.74 to 0.44 due to outliers, whereas the linear correlation on the rank scale has a relatively small effect (0.72 to 0.62). **e) RoDiCE is a copula-based two-sample test.** RoDiCE is an efficient method for testing differences in copula functions between two groups. Rather than a summary measure, such as correlation coefficients, we compare copula functions expressing overall dependence between groups. This allows us to robustly identify differences in complex co-expression structures between two groups of protein complexes to outliers

### 2.1 RoDiCE model

Suppose there are *n* samples, and *g*(*g* = *g*_1_, *g*_2_) represents each condition. We compare two conditions and assume that *g*_1_ and *g*_2_ represent the normal group and the tumor group, respectively. Let **X**_*g*_ = (*X*_1*g*_, *X*_2*g*_,…, *X_Pg_*) be abundances of *P* subunits in group *g*. Given a protein complex, we represent the entire behavior of subunits with a joint distribution **X**_*g*_ ~ *H_g_*(*x*_1_, *x*_2_,…, *x_P_*). The distribution function *H_g_* describes two pieces of information: subunit expression levels and the structure of co-expression between subunits. Meanwhile, copula *C_g_* is a function that can decompose this information into a form that can be handled separately, as follows:

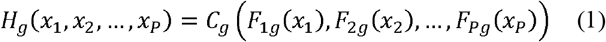

The behavior of each subunit *F_pg_*(*x_p_*) is represented by a distribution function. The copula function itself is a multivariate distribution with uniform marginal probability distribution. The copula function includes all dependency information among the subunits (Nelsen, 2010; Rémillard and Scaillet, 2009; Seo, 2020).

We then use the empirical copula to non-parametrically estimate the copula *C_g_* as it can be widely applicable to various situations and can be represented using pseudo-copula samples defined using rank-transformed subunit abundance 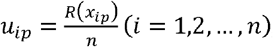;

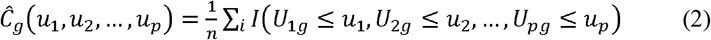

where *R*(·) is a rank-transform function, and pseudo-sample variables are transformed as *R*(**X**_*g*1_) = **U**_*g*1_ and *R*(**X**_*g*2_) = **U**_*g*2_. The empirical copula is robust to noise; however, it represents co-expression structures based on rank-transformed subunit expression levels, which is the so-called scale invariant property in the context of the copula theory (Nelsen, 2010).

To perform DC analysis between groups *g*_1_ and *g*_2_, we consider the following statistical hypothesis:

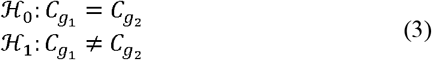

We then derive the following Cramér-von Mises type test statistic to perform statistical hypothesis testing (Rémillard and Scaillet, 2009):

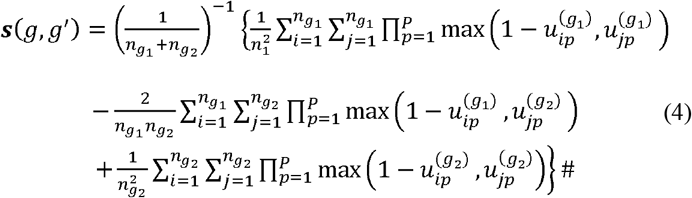

where 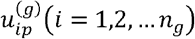 represents the pseudo-observation in group *g*. Note that the computational cost is *n*^2^, where *n*^2^ ≤ *n*_*g*_1__*n*_*g*_2__; *n* = min(*n*_*g*_1__, *n*_*g*_2__). To assess the test statistic (4), we also derived the *p*-value using an algorithm based on Monte Carlo calculations (Rémillard and Scaillet, 2009); however, the computational complexity of the algorithm impedes its application to proteome-wide co-expression differential analysis (results of the simulation experiments described below).

### 2.2 Derivation of statistical significance

Using a permutation test, we derived the *p*-value using the following steps:

1. Randomized concatenated variable from the two groups; **W** = (**U**_*g*_1__, **U**_*g*_2__)
2. Constructed a new randomized variable 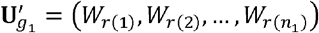 and 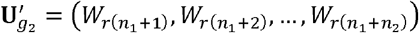 with randomized index *r*(*i*).
3. Replaced copula functions *C*_*g*_1__, and *C*_*g*_2__ in (3) with re-estimated empirical copula function 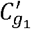 and 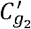 from the randomized samples 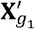 and 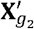.
4. Derived test statistics ***s**′* (*g*_1_, *g*_2_) based on (4) with 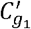 and 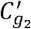

Steps 2 and 3 are indispensable for deriving the null distribution correctly as its derivation by randomization **W**′ = (**U**_*g*_1__, **U**_*g*_2__) alone will distort the distribution, making it impossible to accurately control for type I error (Seo, 2020).

### 2.3 Approximation of *p*-value

The empirical *p*-value is derived as follows:

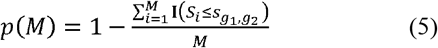

where *M* is the number of randomization and *S_i_* is the test statistic from the null distribution of the *i*-th (*i* = 1, 2,…, *M*) randomization trials. The *p*-value accuracy in (5) is bounded by *p*(*M*) ≥ 1/*M*. As mentioned, calculating the test statistic requires a computational cost of *O*(*n*^2^); therefore, an efficient computational algorithm is required to derive accurate *p*-values in data with a large number of samples. For instance, proteomic cohort projects such as CPTAC/TCGA have more than *n* = 100 samples. To address this problem, we introduced an approximation algorithm for *p*-values based on the extreme value theory (Knijnenburg et al., 2009) and devised a method to calculate accurate *p*-values even with a small number of trials.

The test statistic that exceeds the range of accuracy with randomization trials *M* is regarded as an “extreme value,” and its tail of the distribution could be estimated via a generalized Pareto distribution (GPD), as follows:

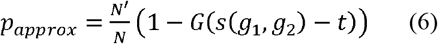

where *N′* is the number of randomized test statistics exceeding the threshold *t* that must be estimated via a goodness-of-fit (GoF) test (Knijnenburg et al., 2009) and *G* is the cumulative distribution function of the generalized Pareto distribution, 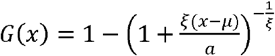 for *k* ≠ 0 and 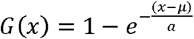 for *k* = 0. To estimate the threshold *t* in (6), the GoF test determines whether the excess comes from the distribution *G*(*x*) via bootstrap based maximum likelihood estimator (Villaseñor-Alva and González-Estrada, 2009). As we do not know a priori, the number of samples sufficient to estimate the underlying GPD with threshold *t*, we must decide the initial number of samples to use. We begin with a large number of samples and increase until the GoF test is not rejected, according to Knijnenburg et al. (2009). As initial samples, we start with those above the 80% quantiles and decrease samples by 1% while the GoF test is rejected. The difference in sensitivity according to the difference in threshold t was also examined in the simulation, however, the choice of threshold did not affect the sensitivity (**Supplementary Data Figure S3**).

### 2.4 Identification of protein complex alteration

As protein complexes show co-expression among multiple subunits (Kerrigan et al., 2011), we hypothesized that the difference in the co-expression structure of the tumor group compared to the normal group is a characteristic quantity of the protein complex abnormality. In previous cancer transcriptome studies, differential co-expression analysis revealed abnormalities associated with protein complexes (Amar et al., 2013; Srihari et al., 2014).

RoDiCE is a flexible model that can robustly capture changes in various co-expressed structural patterns, thus, we must consider what type of structural changes we want to capture. We consider two approaches for the identification of abnormalities in protein complexes. One is that the co-expression structure changes among at least one subunit, and the other is that the overall co-expression structure changes. The former is useful when interested in structural changes in local co-expression and can be used to search for specific targets. The latter may be effective when searching for complexes in which the co-expression structure changes globally.

To capture changes in the co-expression structure among at least one subunit, RoDiCE can be applied with *p* = 2 to robustly capture more than co-expression. In contrast, for global co-expression structural changes (*p* > 2), many combinations of patterns are possible. Therefore, defining in advance what kind of contrasting structures we want to capture will facilitate the interpretation of the results. In this study, we consider the case where the complete co-expression structure (called the “core complex” in Ryan et al., 2017) changes significantly, and we capture the state where the structure is lost in many subunits from the co-expression structure where all subunits are significantly correlated.

In the data analysis of this study, the Spearman’s correlation test was performed on all pairs of subunits within each complex to define the complete correlation structure, and only those complexes that were complete graphs when significantly correlated pairs were set to 1 and others to 0, were analyzed.

### 2.5 Protein membership with protein complex

As we do not know which proteins belong to which protein complexes, we must predict the membership, which can be achieved via two main approaches. One is membership prediction focusing on the modular structure in PPI networks (Adamcsek et al., 2006; Nepusz et al., 2012) and the other is a knowledge-based method using a curated database. We adopted the latter approach, which is based on already validated protein complex membership information, using CORUM (ver. 3.0) (Giurgiu et al., 2019) as a database (see the Supplementary Data for details).

### 2.6 R implementation with multi-thread parallelization

To further accelerate the computation of the test statistic (4) in the randomization steps, we used RcppParallel (Allaire J, 2019). Specifically, we utilized the portable and high-level parallel function “parallelFor,” which uses Intel TBB of the C++ library as a backend on systems that support it and TinyThread on other platforms.

### 2.7 Copula-based simulation model for protein co-expression

Here we provide the outline of a method for simulating co-expressed structures. First, we simulated protein expression levels that showed differential co-expression patterns with outliers in the tumor group and the normal group. We represented the co-expression structure by the covariance parameter in the following multivariate Gaussian copula:

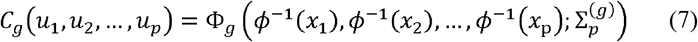

where Φ_*g*_ is the *p* dimensional Gaussian distribution parameterized by *p* × *p* covariance matrix (or correlation matrix) in group *g*, denoted as 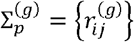, and *ϕ*(*x_i_*) is a univariate distribution. Using the model, we generated the dependency structure with two groups: one with high and the other with low correlations; for the bivariate case (*p* = 2), 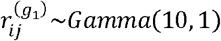 and 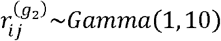, and for the multivariate case (*p* > 2), 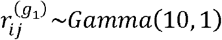 and 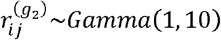. We then generated the co-expression structure using a Gaussian copula with *ϕ*(*x*) = *N*(0, 1) and obtained protein expressions via

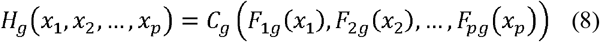

where we set *F_ig_~N*(*μ, σ*) for *i* = 1, 2,…, *p* and *g* = *g* = *g*_1_, *g*_2_ with *μ~N*(2, 1) and *σ~gamma*(2, 1). Furthermore, we added outliers that could affect the co-expression structure. Using the model in (6) and (7), we set the outlier population in both groups as 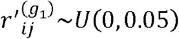 and 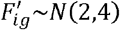 for *i* = 1, 2 and *g* = *g*_1_, *g*_2_ (**Supplementary Data Figure S1**).

### 2.8 Generative model for missing values and imputation

In proteomics, two possible mechanisms exist for missing protein expression levels. One is missing completely at random (MCAR), which involves the accumulation of multiple minor stochastic errors. The other is a non-random missing mechanism due to measurement limitations of LC-MS/MS (missing not at random; MNAR). In actual proteomics data, MCAR and MNAR are thought to be combined to cause missingness. In this study, we reproduced these missing mechanisms using a model based on that proposed by Lazar et al. (2016) and applied the three widely used missing value methods: k-nearest neighbor (kNN), singular value decomposition (SVD), and nonlinear iterative partial least squares (Nipals) to reproduce the noise introduced by missing value imputation. We tested the performance of RoDiCE for several cases of missing data using the overall missing rate α(%) and percentage of MNARs β(%) as parameters. The detailed simulation model is described in **Supplementary Data.**

## 3 Results

### 3.1 Benchmarking RoDiCE with simulation dataset

#### 3.1.1 Type I error control

First, to confirm whether RoDiCE could correctly derive the *p*-value, we performed a test on two groups, with no differences in co-expression structure without outliers, and confirmed the null rejection rate. We performed 100 tests with the proposed method and calculated the null rejection rate at the 1%, 5%, and 10% levels of significance. The same simulation was repeated ten times to calculate the standard deviations. The results show that the proposed method can control type I errors (**Table 1**).

**Table 1.** Type I error controls of the proposed method

|  | p = 3 |  | p = 5 |  | p = 10 |  | p = 20 |  | p = 30 |  |
| --- | --- | --- | --- | --- | --- | --- | --- | --- | --- | --- |
|  | mean | SD | mean | SD | mean | SD | mean | SD | mean | SD |
| 1% | 0.8 | 0.79 | 1.0 | 0.82 | 0.7 | 0.95 | 1.3 | 1.34 | 1.3 | 0.95 |
| 5% | 5.0 | 2.62 | 4.0 | 1.83 | 4.5 | 2.37 | 4.9 | 2.18 | 5.9 | 1.60 |
| 10% | 10.5 | 4.55 | 8.9 | 2.92 | 9.2 | 2.53 | 8.9 | 3.00 | 10.8 | 2.30 |

#### 3.1.2 Robustness to outliers in bivariate case

We then simulated a case in which the co-expressed structure between the two groups differed and outliers were included. We examined the sensitivity of the method to identify a broken co-expressed structure in tumor tissue relative to normal tissue. The results for the bivariate case are presented in Figure 3.

**Fig 3.**
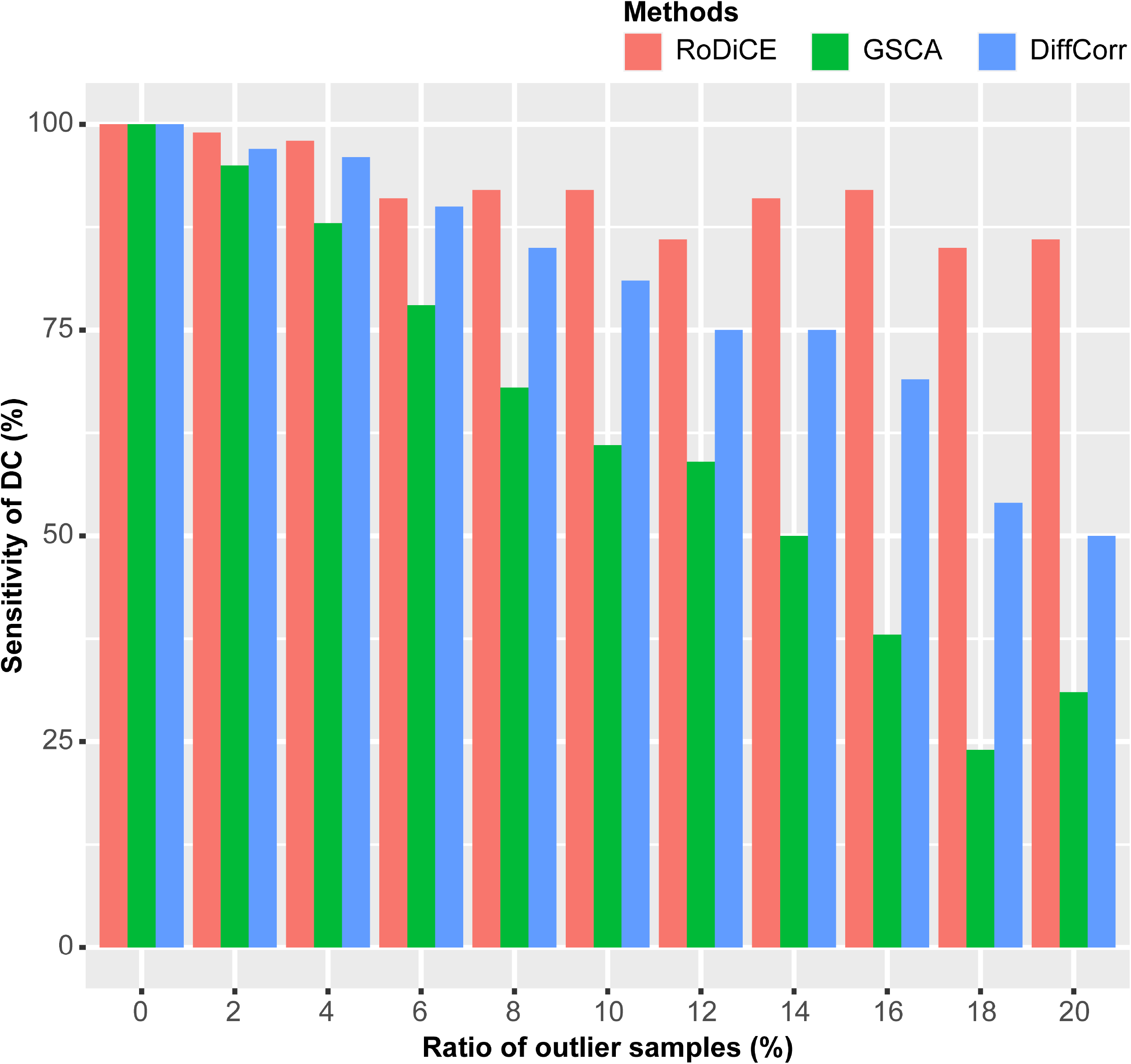
Sensitivities and ratio of outliers (bivariate case). The percentage of outliers is taken on the horizontal axis, and the sensitivity of the co-expression differences by each method (5% level of significance) is shown on the vertical axis.

To demonstrate the advantages of the proposed method, we examined the sensitivity of increasing the percentage of outliers in 2% increments from 0% to 20% and compared it further with DiffCorr and GSCA, a two-group co-expression test method based on Pearson’s linear correlation (**Figure 3**). For outliers, the proposed method showed robust co-expression test results, with an accuracy of more than 85% up to a percentage of outliers of approximately 15%. Conversely, the sensitivity of the method based on linear correlation begins declining from the level of 2% outliers, and for data containing 15% outliers, the sensitivity drops to ~ 30%. In contrast, there was no significant difference in specificity (**Supplementary Data Figure S2**). We also examined the relationship between sample size and sensitivity. RoDiCE showed the highest sensitivity (**Supplementary Data Figure S4)**.

#### 3.1.3 Robustness to outliers in multivariate case

Next, we examined the multivariate case (p ≥ 3) by determining the sensitivity of RoDiCE to co-expression changes when the co-expression structure was altered by one for different dimensions and percentages of outliers. The number of dimensions was set to p = 3, 6, 10, 20, and 30, and the percentage of outliers was set to 0%, 5%, and 10%. The number of samples was set to *n* = 100. GSCA (Choi and Kendziorski, 2009) and GSNCA (Rahmatallah, et al., 2014) were chosen for comparison.

To understand the characteristics of each method, we first examined the sensitivity of simultaneous DC without outliers, i.e., noise-free case (**Top panels in Figure 4**). GSCA tends to capture the local structure as it considers the sum of the DC in each pair of variables. In contrast, RoDiCE tends to capture the broader structure of DCs as it captures the differences in joint distribution, which is also true for GSNCA, which captures the differences by eigenvectors of covariances.

**Fig 4.**
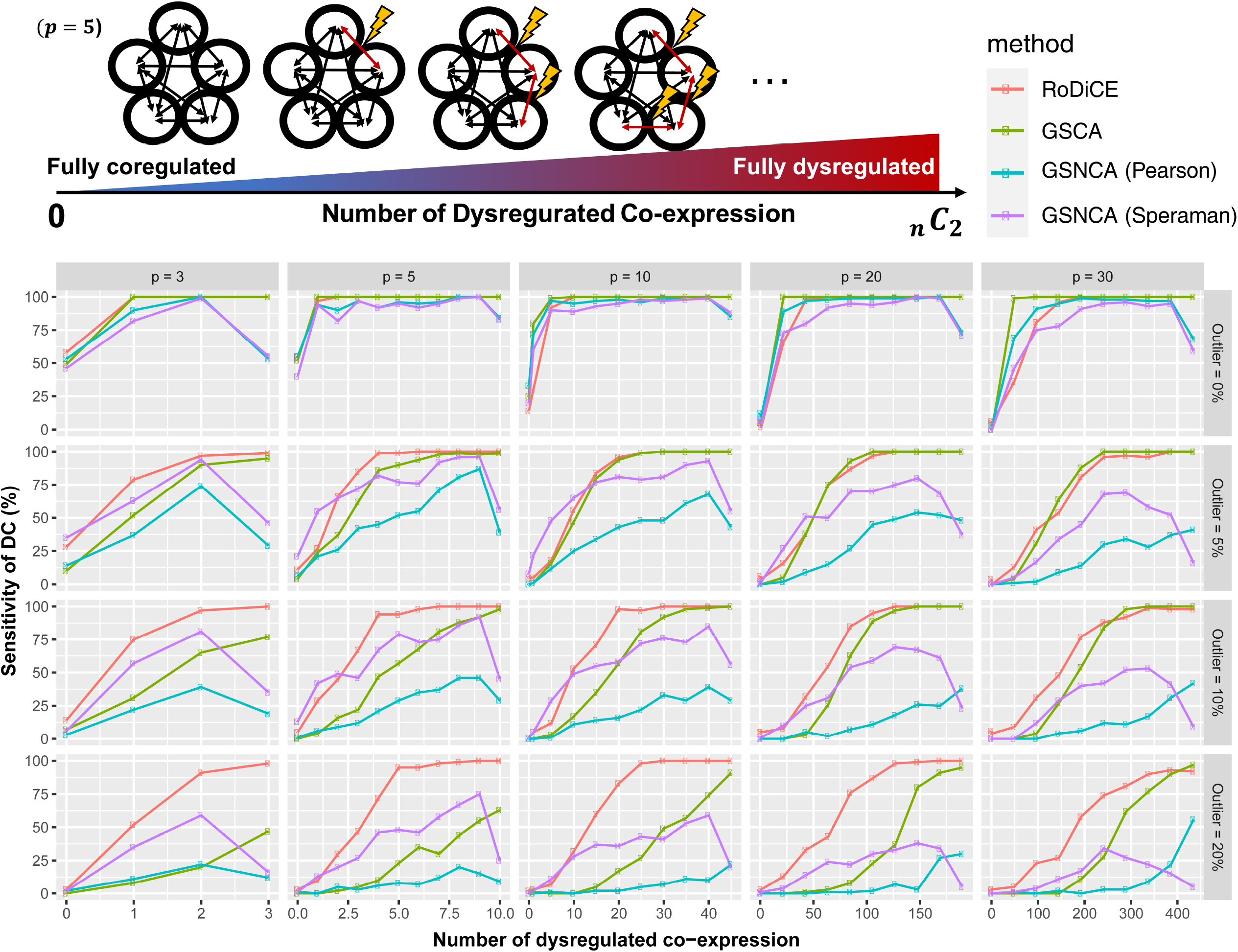
Sensitivities and ratio of outliers (multivariate case). The percentage of outliers is taken on the horizontal axis, and the sensitivity of the co-expression differences by each method (5% level of significance) is shown on the vertical axis.

Next, we investigated the robustness to outliers (**From second to bottom panels in Figure 4**). Considering that the copula converts protein expression into rank space, it is negligibly affected by outliers. Therefore, there was no significant difference between the case of 0% outliers and the cases of 5% and 10% outliers in any number of dimensions. In contrast, the other methods showed an overall lower sensitivity. In particular, GSCA, which was sensitive to changes in the local co-expression structure, was rarely captured by the outliers compared to the noise-free case.

#### 3.1.4 Robustness to noise generated by missing value imputation

In addition, we show the performance of the proposed method to the noise generated during the imputation of missing values. The sensitivity and specificity are shown in **Figure 5 (panels in the top two rows)** for different combinations of (*α, β*). For sensitivity, GSCA was superior in all dimensions, followed by RoDiCE. In contrast, for specificity, RoDiCE and GSNCA were superior and GSCA had the lowest specificity results. Based on these observations, we evaluated the balanced performance in terms of both sensitivity and specificity, with a likelihood-like index, L, as follows:

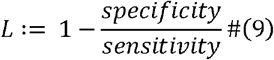

and compared the DC methods (**Figure 5**). First, it can be seen that RoDiCE has generally smaller variation in *L* than the other methods for different missing patterns in terms of different combinations of (*α, β*) over the missing imputation methods and dimensions. This implies that the proposed method does not relies heavily on the DCs specifically produced by the missing rate and missing value mechanisms, suggesting that it may be able to maintain stable performance for different and various data sets. GSNCA also seems to have a similar performance after RoDiCE, however, the variation is relatively large and the actual values of sensitivity and specificity are lower than those of RoDiCE, in most cases (**The panels in the top two rows of Figure 5**). GSCA showed a tendency for *L* to be greater than 1 compared to the others, resulting in a high false negative rate.

**Fig 5.**
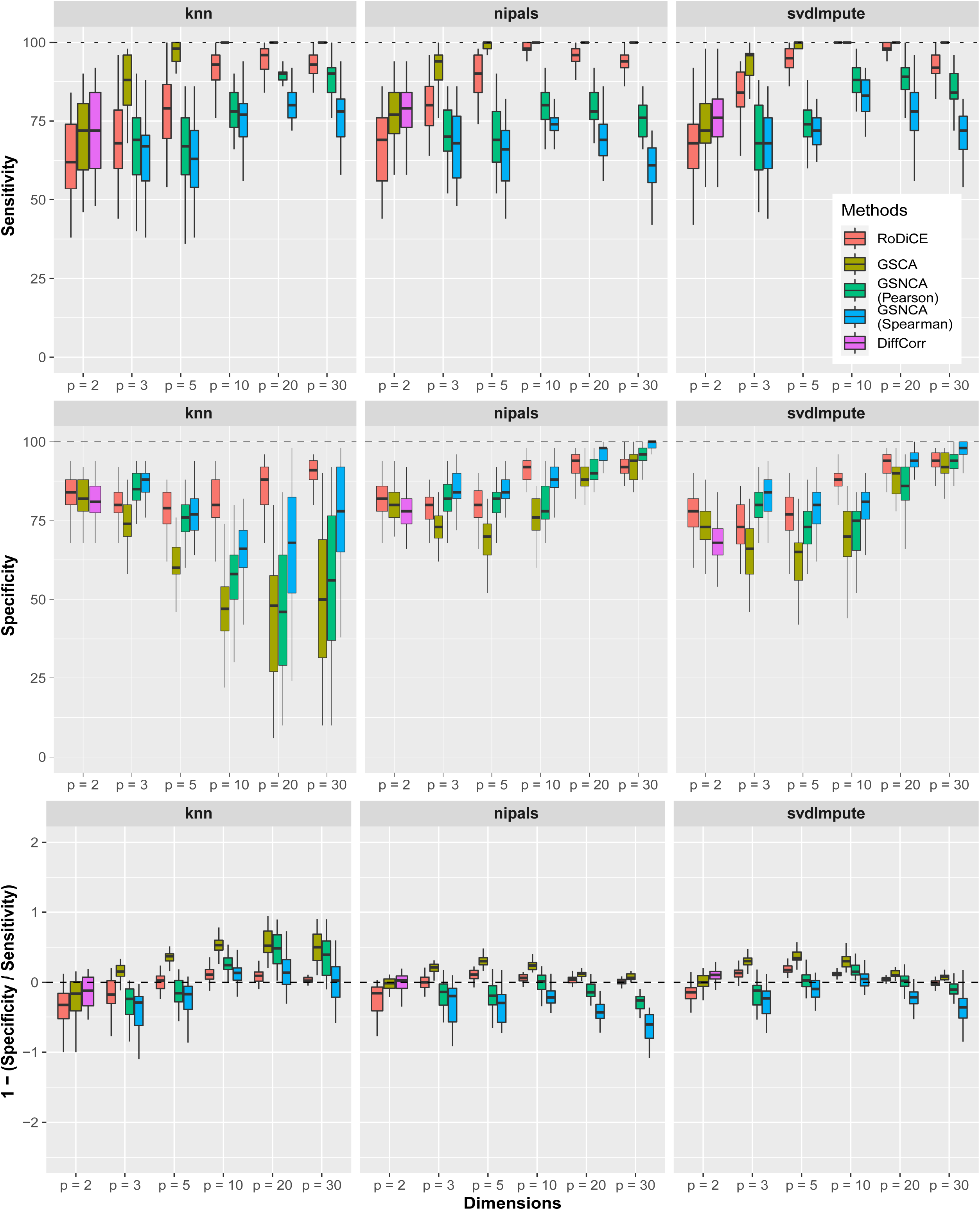
Comparing performance of DC methods for missing pattern and imputation methods. The boxplots show the distribution of sensitivity (%) (upper panel), specificity (%) (middle panel), and L (lower panel) derived by each DC method after imputing missing data generated by a combination of control parameters (α,β), for each missing value imputation method (knn for kNN, nipals for Nipals, and SVD for svdImpute). In the lower panel, L=0 (dashed line) indicates that the false positives and false negatives are controlled to the same extent, while L > 0 indicates that the false positives tend to be high. L > 0 indicates that the false positive rate tends to be high. Conversely, L < 0 indicates that the false negative rate tends to be high.

In the comparison among the missing value imputation methods, Nipals and SVD, based on principal component analysis, which is a missing value imputation method considering the covariance structure, showed better overall performance than kNN in terms of sensitivity, specificity, and *L*s.

### 3.2 Computational performance

Finally, we also examined the computational speed, comparing it with the R package TwoCop, which implements the Monte Carlo-based method (Rémillard and Scaillet, 2009) used for the two-group comparison of copulas (**Table 2**). The proposed method is ~70 times faster than TwoCop for two variables and more than 5000 times faster for 30 variables, while having the same accuracy as TwoCop (**Supplementary Data Figure S5**), and is sufficiently efficient as a copula-based two-group comparison test method (**Table 2**). In contrast, the estimation of the copula function required more computational time than the linear correlation coefficient-based method due to the computational complexity of estimating the copula function.

**Table 2.** Computation time for ten replicates

|  | p = 2 | p = 3 | p = 5 | p = 10 | p = 20 | p = 30 |
| --- | --- | --- | --- | --- | --- | --- |
| DiffCorr | 0.001 | * | * | * | * | * |
| GSCA | 0.152 | 0.096 | 0.096 | 0.096 | 0.096 | 0.096 |
| GSNCA | * | 0.841 | 1.049 | 1.496 | 2.405 | 5.584 |
| RoDiCE | 0.373 | 0.457 | 0.499 | 0.605 | 0.771 | 0.986 |
| TwoCop | 25.301 | 51.482 | 125.753 | 437.747 | 1674.990 | 5101.482 |

### 3.2 Application to cancer complexome analysis

To provide a real-world example of the applications for RoDICE, here, we assess clear renal cell carcinoma (ccRCC) data published by CPTAC/TCGA (Clark et al., 2019) with RoDiCE. The data are available from the CPTAC data portal (https://cptac-data-portal.georgetown.edu) in the CPTAC Clear Cell Renal Cell Carcinoma discovery study. The data labeled “CPTAC_CompRef_CCRCC_Proteome_CDAP_Protein_Report.r1” were used. In the following analysis, only protein expression data that overlaps with protein groups in human protein complexes in CORUM and in CPTAC were used. Missing values were completed based on principal component analysis, and the missing values were completed by ten principal components using the pca function in pcaMethods.

For the complete data, RoDiCE was applied to the normal and cancer groups for each protein complex. FDR was calculated by correcting the p-value for each complex using the Benjamini–Hochberg method.

#### 3.2.1 Anomalous complexes detected by pairwise comparisons

We identified anomalous protein complexes in protein expression data from 110 tumor and 84 normal samples; out of 3,364 protein complexes in CORUM, 1,244 (7,937 out of 57,980 protein pairs) contained at least one co-expression difference between subunits with FDR ≤ 5% (**Supplementary Data, Table S1)**. DiffCorr, which showed the second-best performance in numerical experiments, identified differential co-expression in 11,699 pairs, 3278 of which were common to RoDiCE (**Supplementary Data Fig. S6**). Although the number of differential co-expression identified by RoDiCE was conservative, it is a reasonable result considering RoDiCE is robust in terms of sensitivity and specificity to noise due to outliers and missing value imputations.

The proposed method identified several protein complexes containing driver genes on regulatory signaling pathways in ccRCC (**Figure 6a**) (Li et al., 2019). The identified pathways included known regulatory pathways important for cancer establishment and progression, starting with chromosome 3p loss, regulation of the cellular oxygen environment (VHL), chromatin remodeling, and disruption of DNA methylation mechanisms (PBRM1, BAP1). They also included abnormalities in regulatory signals involved in cancer progression (AKT1). Moreover, several identified complexes included key proteins, for example, MET, HGF, and FGFR, which could be directly inhibited by targeted drugs, such as Cabozantinib and Lenvatinib. Considering that a previous study reported that sensitivity to knockdown several genes was well associated with expression levels of protein complexes (Nusinow et al., 2020), co-expression information on protein complexes containing druggable genes might be useful for optimizing drug selection.

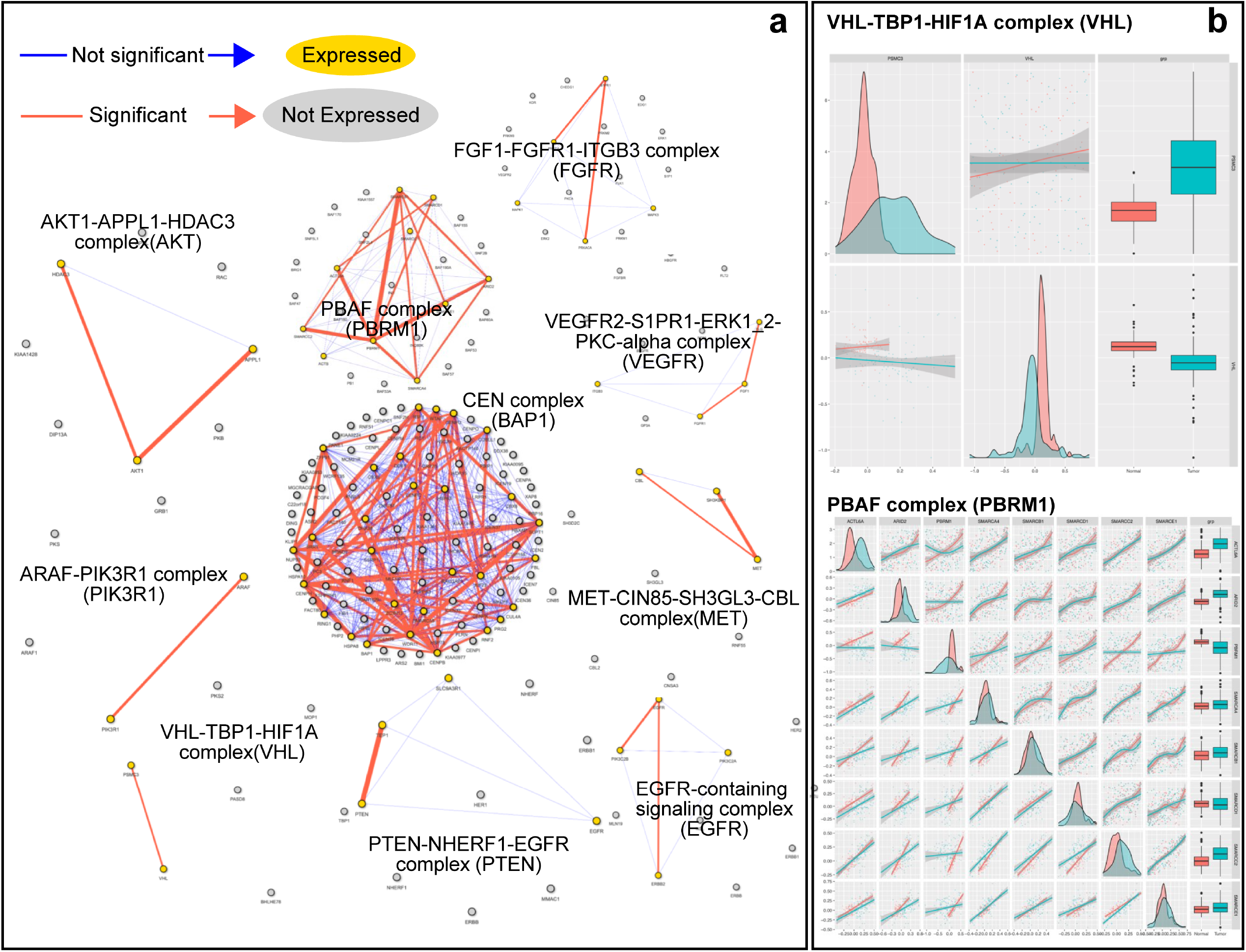

A close examination of the above-identified protein complexes allows us to partially understand how the dysregulation of protein was a co-expression abnormality between VHL and TBP1. The upregulation of TBP1 is known to induce dysregulation of downstream HIF1A molecules in a VHL-dependent manner (Corn et al., 2003). In fact, the protein expression of TBP1 increased in the tumor group. We also examined the PBAF complex containing the driver gene *PBRM1*, which is thought to occur following VHL abnormalities. Along with a decrease in PBRM1 protein expression, there was a loss of tumor group-specific co-expression structure among many subunits involved with PBRM1 levels.

#### 3.2.2 Anomalous complexes detected by groupwise

To identify abnormal cases in the complete co-expression structure, we applied RoDiCE to the joint distribution of inter-subunit co-expression with *p* > 2. First, 394 of the CORUM-registered complexes showed complete co-expression structures in the normal group (p-value ≤ 5% for each pair of subunits by pairwise Spearman correlation test). The largest complex was the respiratory chain complex I (holoenzyme) with 41 subunits (*p* = 41), followed by the 28S ribosomal subunit (*p* =30), which was identified as a complex with a complete co-expression structure (**Supplementary Data Table S2**). Most complexes consisted of three to four subunits.

Of these, 136 complexes with FDR≤ 5% were identified by RoDiCE as having abnormal co-expression structures (**Supplementary Data, Table S2, Figure. S7-S9**). The large complexes (*p* ≥ 10) with differential co-expression included respiratory chain complex I, 28S ribosomal subunit, and CCC-Wash. Among the medium-sized protein complexes (5 ≤ *p* < 10) were cytochrome c oxidase, mitochondrial proteins, the conserved oligomeric Golgi complex (COG), and TNF-α/NF-κB signaling complex. The smaller protein complexes (3 ≤ *p* < 5) were the DNA repair complex NEIL1-PNK-Pol(β)-LigIII(α)-XRCC1, SNX complex, SNARE complex, and PDGFRA-PLC-γ-1-PI3K SHP-2 complex.

Some complexes have been reported as cancer-specific abnormalities, including those associated with renal cancer. For example, ribosome complexes have a low correlation with mRNA and protein by proteogenomic analysis, suggesting a protein level specific regulatory mechanism that may be an important therapeutic target in renal cancer, although the detailed mechanism remains unclear (Cleark et al., 2019, Devlin et al., 2016). Respiratory chain complex I is a tumor suppressor (Lemarie et al., 2011), and numerous cancer-specific mutations in its subunits have been reported. In this study, we found that the co-expression structure between many subunits was highly abnormal. At the same time, the protein expression of mitochondrial subunits was also down-regulated (**Supplementary Data Figure S9**). Mitochondrial subunits are known to undergo copy number alterations and reduced mRNA expression in many cancer types, and our analysis is consistent with these findings at the protein level (Reznik, et al., 2017).

## 4 Discussion

In this study, we developed an algorithm of robust identification for protein complex aberrations based on differential co-expression structure using protein abundance. Protein expression data measured through LC-MS/MS contains a non-negligible percentage of outliers due to technical limitations and variation due to biological reasons such as post-translational modifications and missing values. This causes over- (or under-) estimation of co-expression. However, the copula-based DC approach is a powerful statistical framework that serves as a solution to this problem.

To the best of our knowledge, statistical models that consider noise introduced by post-translational modifications and missing values have not been sufficiently studied in proteome analysis. In particular, in the presence of missing values, it is largely unknown how much distortion of the original co-expression structure is caused by missing value imputation, and how it may affect functional protein network analysis and differential co-expression. Further comprehensive systematic investigations will be indispensable for the establishment of large-scale proteome analysis methods in the future.

In addition to noise robustness of the proposed method, another key property of the copula that is important for capturing the co-expression structure, is self-equitability (Chang et al., 2016; Ding et al., 2017). Copulas can capture nonlinear structures between variables, and self-equitability allows for evaluation of the degree of dependency equally between variables in linear and nonlinear relations. Therefore, copula allows us to compare a much broader range of co-expressed structures, compared to conventional linear and nonlinear correlations.

The copula-based co-expression analysis approach is a powerful modeling method for data sets with expected noise, although there remain challenges in high-dimensional estimation. In particular, it could be useful for modeling proteome-wide protein expression patterns. The proposed approach is useful for understanding abnormalities in the protein complexes of cancer. However, studies focusing on protein complexes in large-scale cancer proteomics are in their infancy. We believe that this approach will, therefore, provide valuable insights into the molecular mechanisms of cancer and the search for new drug targets.

