## Supplementary material for "RoDiCE: Robust differential protein co-expression analysis for cancer complexome": Supp_manuscript_r2.pdf

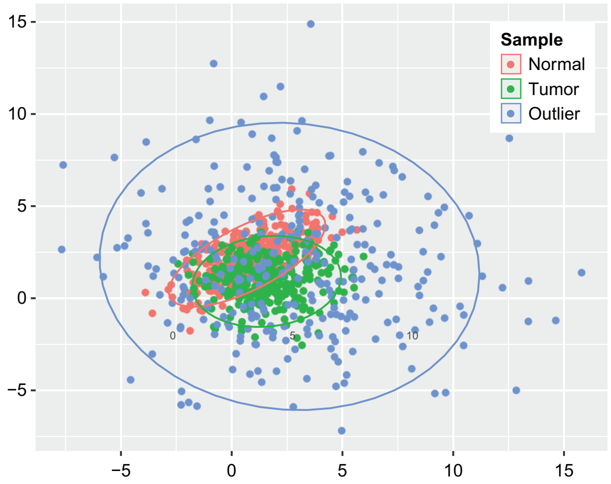


**Fig S1. Simulated dataset.** Generated samples in the numerical experiments for the bivariate case. To mimic the noise in the proteome abundance dataset, the outlier population was assumed other than that of the tumor and normal population.

### 1. Details of the generative model for missing values and imputation

The main parameters of the missing mechanism are $\alpha$ (%), the total percentage of missing values, and $\beta$(%), the percentage of missing values accounted for by MNARs. First, let the measurement limit $T$ be generated by $T \sim N(q, 0.01)$ using the $\alpha$(%) quantile $q$ of all measurements. If the observed values are $< T$, the $\beta\alpha/100$ (%) of the observed values is missing according to the Bernoulli trial $Ber(\beta\alpha/100)$, and the remaining $(100-\beta\alpha)/100$ (%) of the missing values is introduced randomly.

In this simulation, we generated a Gaussian copula according to Equations (7) and (8) for $p = 100$, $r_{ij}^{(g_1)} \sim Gamma(5, 1)$ and $r_{ij}^{(g_2)} \sim Gamma(1, 5)$ and a joint distribution, the marginal distribution of which consists of $\mu \sim N(2,1)$ and $\sigma \sim gamma(2,1)$. When generating co-expressed structures by the Gaussian copula, we assumed protein complexes of multiple sizes and set up block structures that were co-expressed with each other (block structures with $p = 2,3,5,10,20,30$ were always included, and the rest were generated with random sizes). We introduced missing values for the obtained protein expression levels and generated missing patterns in 10% increments for $\alpha \in [0,40]$, $\beta \in [0,80]$ for multiple scenarios.

For kNN, we used the impute function implemented in the R package MSnBase to allow up to 90% missing values in each row and column and used default parameters for the rest. For Nipals and SVD, the pca function of the R package pcaMethods was used and method parameters were set to “nipals” and “svdImpute”. The number of principal components, called nPCS argument, was set to 10, the upper limit of the number of iterations was set to 10,000, and the default parameters were used for the rest.

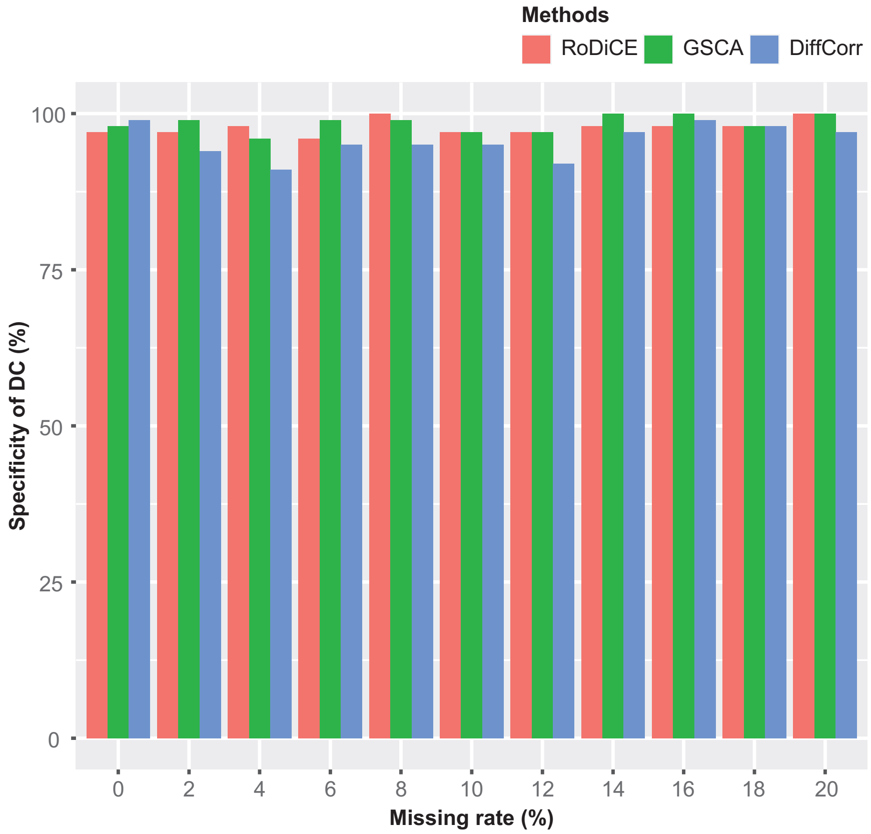


**Fig S2. Specificity and ratio of outliers (bivariate case).** The percentage of outliers is presented on the x-axis, and the sensitivity of the co-expression differences by each method (5% level of significance) is shown on the y-axis.

### 2. Thresholds of GPD and DC sensitivity

To approximate the p-value using GPD, it is necessary to determine the threshold t in Eq. (6). Although there is no general method to determine the threshold t, it is preferable to use as many samples as possible as the hem of the distribution is to be extrapolated [2]. Here, we provide a guideline based on a simulation experiment, in which we use the same model as that used when we conducted the multivariate accuracy verification. Considering that the purpose of this study is to examine the approximate accuracy of $p$-values, outliers were excluded.

The number of samples was set to $n = 50$ and 100, and in each case, $p(M)$ in Eq. (5) obtained by permuting $M = 1000$ and $M = 10{,}000$ times was approximated by GPD with a smaller number of permutations ($M' = 100, 200, 400, 600, 800, 1000$). The accuracy of the approximation was examined by comparing the $p_{approx}$ obtained by GPD approximation with a smaller number of permutations ($M' = 100, 200, 400, 600, 800, 1000$). The accuracy was evaluated by comparing the p_approx obtained by approximating the GPD with M^'=100, 200, 400, 600, 800, and 1000, where the significance level was set at $5\%$. In other words, given indicator function I(·), the accuracy is given as

$$r = \frac{\sum_{i=1}^{m} I\left(p_{i,approx} \leq 0.05\right)}{\sum_{i=1}^{m} I(p_i(M) \leq 0.05)}$$

where $m$ is the number of simulations and $m = 100$ in this case. The simulations were performed for different numbers of dimensions $p = 3, 5, 10, 20, 30$ and threshold values $t = 0.4, 0.5, 0.6, 0.7, 0.8, 0.9$, the results for which are shown in Figure S3. There was no significant difference in the results based on the choice of threshold $t$. In contrast, when the number of permutations $M'$ was < 200, the approximation accuracy was lower than the other M' values. In addition, the approximation accuracy does not depend on the number of dimensions p. Thus, we recommend the number of permutations to be at least ~300 times to approximate the number of permutations of $10^3$.

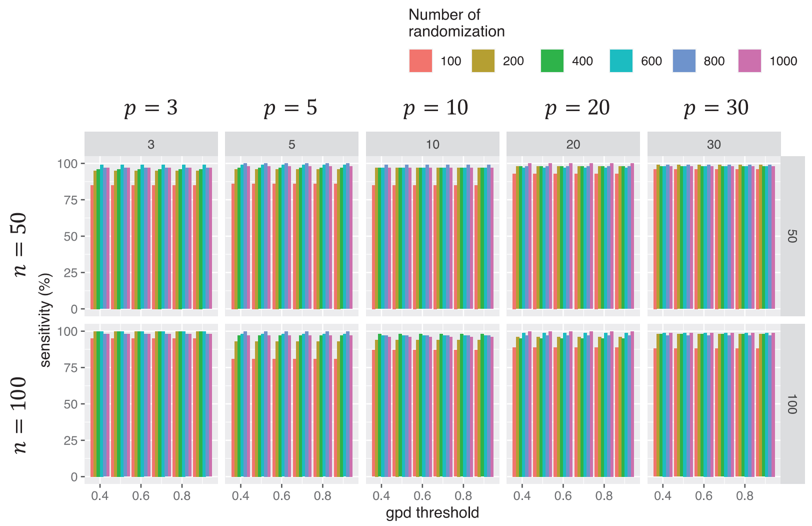


**Fig S3. Sensitivity and GPD threshold**

### 3. Sample size and accuracy

To investigate the relationship between sample size and identification accuracy, we simulated the sensitivity of RoDiCE while increasing the number of samples in increments of 10 from 30 to 100 samples. All other settings were the same as those in **Figure 3**, with the exception of outlier percentage, which was set at 5%.

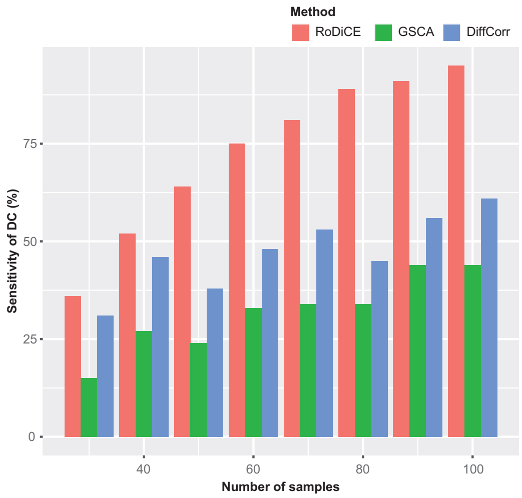


**Fig S4. Sensitivities and sample size.** The x-axis shows the sample size, whereas the y-axis shows the sensitivity of the co-expression differences by each method (5% level of significance).

**4. Comparison of p-values between RoDiCE and TwoCop**

We compared the p-values computed by RoDiCE and TwoCop, and based on the simulation model in Section 2.7, we generated a multidimensional Gaussian distribution with p = 3 and n = 100. The co-expression structure in each group was generated randomly and the p-values in various cases were calculated; $r_{ij}^{(g_1)} \sim U(1,10)$, $r_{ij}^{(g_2)} \sim U(1,10)$. We calculated the p-values for 300, 500, and 1000 permutations. The number of simulations was set to 100 for each case.

Fig. S4 shows the scatter plots of p-values for RoDiCE and TwoCop for each number of permutations. For any number of permutations, The correlation coefficients of the p-values obtained in each of the 100 simulation trials were near 1, indicating that RoDiCE and TwoCop have the same accuracy.

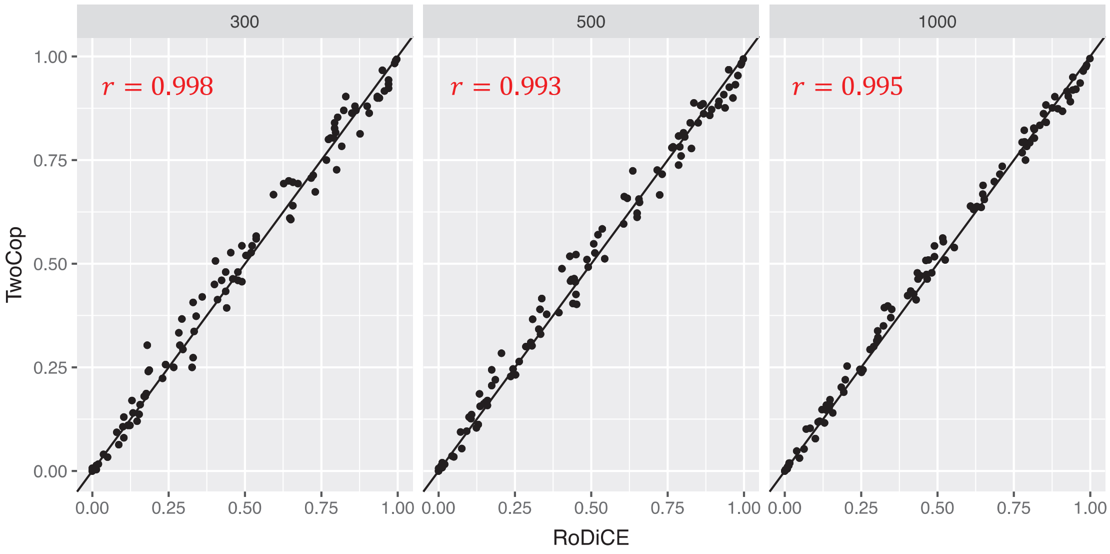


**Fig S5.** Comparison of p-values between RoDiCE and TwoCop

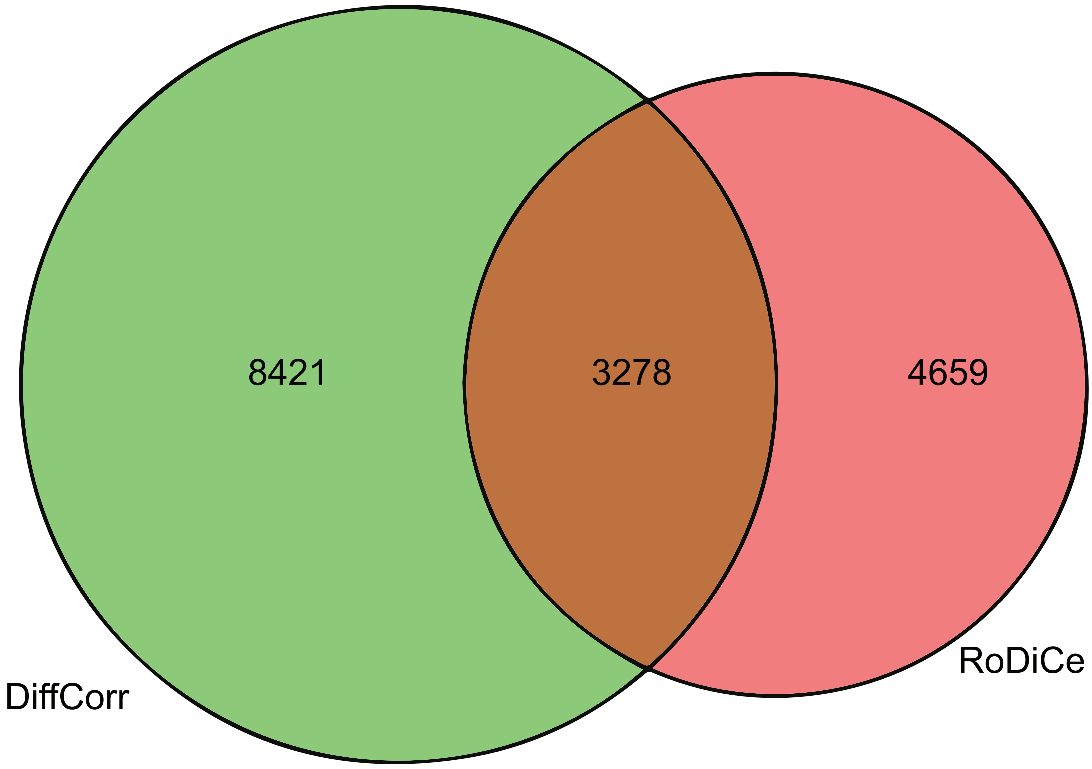


**Fig S6. Total number of abnormal complexes identified by pairwise comparison**

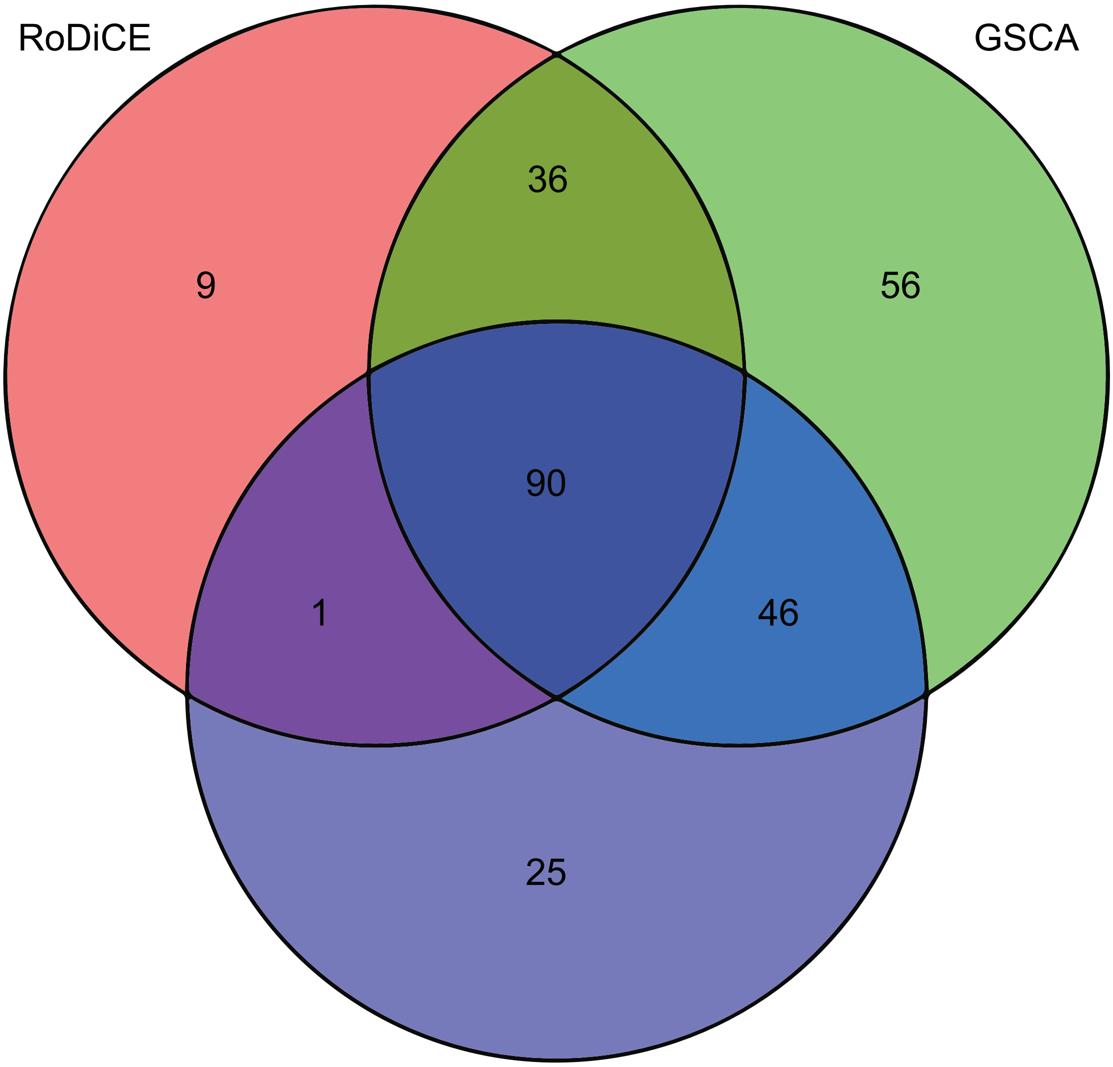


**Fig S7. Total number of abnormal complexes identified by groupwise comparison.**

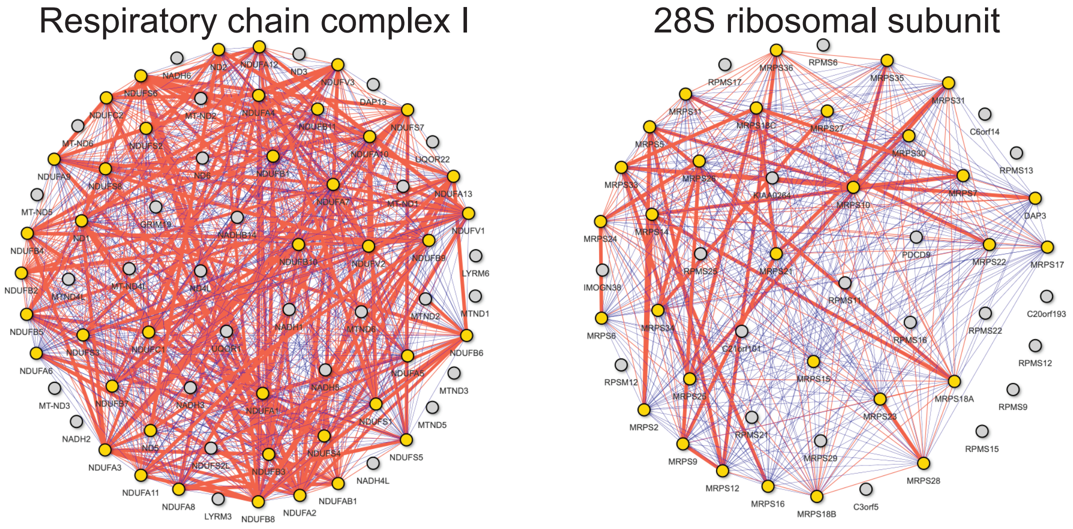


**Fig S8. DC network of the respiratory chain complex I and 28S ribosomal subunit.**

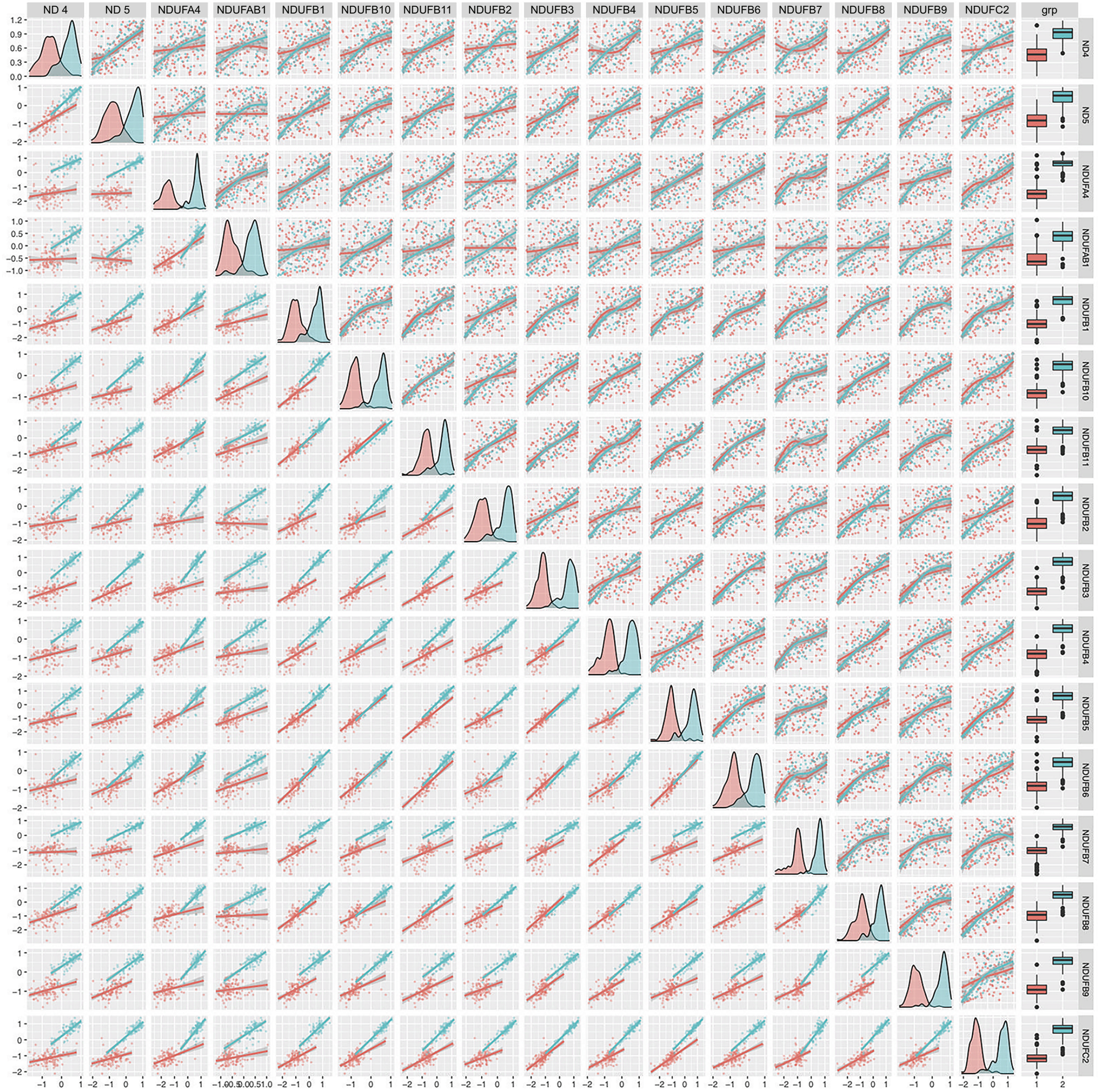


**Fig S9. Co-expression of the respiratory chain complex I with the two groups**
